# Nanotopography reveals metabolites that maintain the immunosuppressive phenotype of mesenchymal stem cells

**DOI:** 10.1101/603332

**Authors:** Ewan A Ross, Lesley-Anne Turner, Anwar Saeed, Karl V Burgess, Gavin Blackburn, Paul Reynolds, Julia A Wells, Joanne Mountford, Nikolaj Gadegaard, Manuel Salmeron-Sanchez, Richard OC Oreffo, Matthew J Dalby

## Abstract

Mesenchymal stem cells (MSCs) are multipotent stem cells that are immunosuppressive and thus of considerable therapeutic potential in transplant operations. However, MSCs rapidly differentiate once in culture, making their large-scale expansion for use in immunosuppressive therapies challenging. Although the differentiation mechanisms of MSCs have been extensively investigated using materials, little is known about how materials can modulate paracrine activities of MSCs. Here, we show for the first time that nanotopography can control the immunomodulatory capacity of MSCs through decreased intracellular tension increasing oxidative glycolysis. We also use the nanotopography to identify bioactive metabolites that modulate intracellular tension, growth and immunomodulatory phenotype of MSCs in standard culture. Our findings show a novel route to support large-scale expansion of functional MSCs for therapeutic purposes

---

Research has provided scientists with a variety of materials for investigating the physical properties that stimulate mesenchymal stem cell (MSC) differentiation^1–4^. The materials properties that have been exploited include stiffness^1^, adhesivity^2,3^, chemistry^5^ and nanotopography^4^. The use of these tools has revealed that MSC differentiation along the adipogenic pathway is characterised by cells with low adhesion, low cytoskeletal tension, and by the suppression of the bone-related transcription factor, Runt-related transcription factor 2 (RUNX2)^2,3,6^. By contrast, MSC osteogenic differentiation is characterised by cells with high levels of adhesion and intracellular tension, mediated by Rho-A kinase (ROCK), as well as by RUNX2 activation^2,3,6^. Materials control of MSC differentiation will be of great value in biomaterials and tissue engineering approaches.

The potential therapeutic applications of MSCs are now extending beyond tissue engineering, towards use of MSCs as cellular therapies. To support this growing demand, it is critical that we develop methods to grow sufficient numbers of high quality, stable MSCs. This is important because *ex vivo,* out of their niche, MSCs rapidly and spontaneously differentiate, mainly because they are primed to differentiate by the unfamiliar stiffness and chemistry of polystyrene surfaces within culture flasks, which were developed for phenotypically stable cells^7^.

The major focus for MSC use as a biological therapeutic is in suppression of immune response. Via paracrine signalling, MSCs can reduce the immune response and inflammation^8^ and this capability is currently under evaluation in human trials for graft vs host disease following haematopoietic stem cell transplantation^9^, and in the co-transplantation of MSCs with islet cells in the treatment of diabetes^10,11^. However, it is important to note that large-scale MSC production, which is required to generate sufficient quantities of clinical grade cells, is offset by a reduction in their immunosuppressive capability, which typically occurs following their long-term culture^12^. Furthermore, the currently advocated MSC expansion protocol relies on the use of multi-layer, large surface-area, cell-culture ware^13^.

A small range of materials that can promote prolonged *in vitro* MSC self-renewal have been identified by utilising nanotopography^14^, surface chemistry^15^, elasticity^16^ and micro-contact printed adhesive islands^17^. These studies demonstrate that when adhesion is reduced, relative to adhesion levels in MSCs cultured on polystyrene, but not reduced to the level that would promote adipogenesis, stem cell multipotency can be retained^16–18^. However, understanding of biological mechanism is nascent and ability to influence MSC immunosuppressive capability is unknown.

In this study, we use defined nanotopographies to investigate if the materials, such as those that have been successfully used to study MSC multipotency and differentiation, can be used to study their immunosuppressive properties as well. We also investigate if the materials that stimulate enhanced MSC immunosuppressive capacity can be used to dissect the molecular mechanisms involved. We hypothesise that such materials could be used to highlight phenotype-specific targets, providing a potential means by which to identify metabolites that might serve as biological small molecules that could help to address the major challenges in the therapeutic manufacture of MSCs.

## Nanotopography can maintain MSC immunosuppressive capacity

Nanotopography has been previously demonstrated to promote MSC multipotency in culture without suppressing cell growth; specifically, nanopits of 120 nm diameter, 100 nm depth and 300 nm centre-to-centre spacing within a square arrangement (SQ, Fig. 1A)^14^. To investigate if this surface can also retain MSC immunosuppressive capacity, a T cell proliferation assay was used. In this assay, human peripheral blood mononuclear cells (PBMCs) were labelled with the intracellular proliferation dye CFSE (carboxyfluorescein succinimidyl ester) and were stimulated with phytohemagglutinin (a mitogenic lectin) and interleukin-2 (IL-2) to drive the proliferation of T cells. CFSE-labelled PBMCs were then co-cultured with primary human MSCs for 5 days. The ability of MSCs to reduce T cell proliferation was measured through the detection of CFSE dilution by flow cytometry (upon cell proliferation, each daughter cell contains half the amount of intracellular CFSE compared to the mother cell; see Fig. S1). Stimulated T cells in the absence of MSCs were used as a positive control and T cells cultured in the absence of any stimulation as a negative control.

**Fig. 1.**
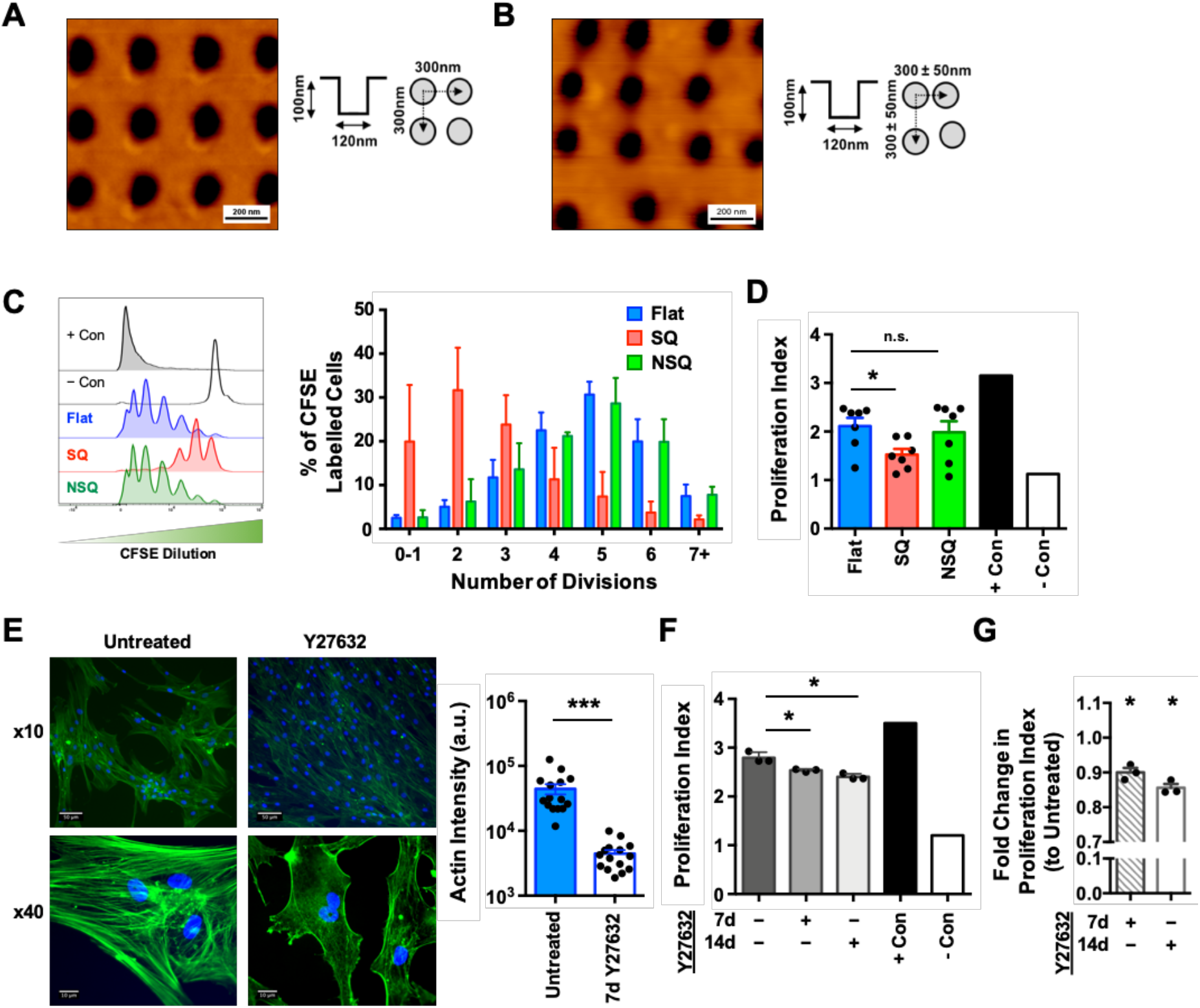
Nanotopography can maintain MSC immunosuppressive capacity via decreased intracellular tension. (**A**) Representative atomic force microscopy images of square (SQ) patterned and (**B**) osteogenic-enhancing, offset near square (NSQ) polycarbonate nanotopographies. (**C**) Stro-1^+^ MSCs were cultured on nanotopographies for 14 days, then cocultured with CFSE-labelled, PHA and IL-2 stimulated PBMCs for a further 5 days. CFSE dilution was quantified by flow cytometry (left panel) and graph shows representative results from one co-culture (n=4 topographies per group, mean ± S.D.). (**D**) Proliferation index was calculated to allow comparison of MSCs immunosuppressive potential from multiple donors (n=7 donors). (**E**) MSCs were cultured on flat nanotopographies for 14 days in the presence or absence of the ROCK inhibitor, Y27632. Actin cytoskeleton changes were revealed by phalloidin staining (n=15 fields per group). (**F**) The immunosuppressive capacity of cells cultured on flat topographies for 7 or 14 days in the presence or absence of Y27632 was assessed as in (**C**) (n=3 topographies per group, mean ± S.D.). (**G**) Fold change in proliferation index to untreated controls of MSCs grown on flat topographies in the presence of Y27632 for 7 or 14 days. Means ± SEM and number of donors (N) are shown for each condition. \*\*\**p* < 0.0001; \**p* < 0.05; n.s., non-significant.

Stro-1^+^ selected, bone marrow-derived, human skeletal MSCs were cultured for 14 days on the SQ nanotopography, which had been injection moulded into polycarbonate. As positive controls, we used a flat topography and an osteogenic-differentiation promoting nanotopography^4^ (both fabricated in polycarbonate). The differentiating nanotopography is similar to SQ but pits are offset by up to ± 50 nm in the x and y axes from the centre position^4^, and is termed near square (NSQ, Fig. 1B).

CFSE-labelled T cells were added to MSCs and co-cultured for five days, followed by flow cytometry analysis. This showed that while MSCs cultured on flat and NSQ nanotopographies displayed poor immunosuppressive capabilities, as measured by increasing numbers of higher-division T cells, T cells cultured with MSCs on the SQ surface underwent fewer divisions (Fig. 1C). This is supported by the reduced proliferative index of T cells co-cultured with MSCs on SQ (Fig. 1D), relative to the other nanotopographies, indicating that the immunosuppressive capacity of MSCs on SQ is significantly increased (p = 0.0245). This is the first time a material alone has been shown to enhance the immunosuppressive capability of MSCs.

It is hypothesised that multipotent MSCs have intermediate levels of intracellular tension, slightly lower than that of fibroblasts^18,19^. As has been described, ROCK is a central mediator of intracellular tension in MSCs^2,3^. We therefore hypothesised that reducing ROCK-mediated intracellular tension in MSCs on flat control surfaces, using the ROCK inhibitor Y27632, should drive these MSCs (which display a fibroblastic phenotypic drift, typical of MSCs in cell culture^14^), to behave more like immunosuppressive MSCs on SQ. Morphological change was indeed observed (Fig. 1E) along with reduction in proliferation index of T cells co-cultured with MSCs on flat control surfaces with ROCK inhibition (Fig. 1, F and G). Thus, MSCs on flat controls with reduced intracellular tension display a more immunosuppressive phenotype. These results support the hypothesis that the MSC phenotype is influenced by intracellular tension, with greater intracellular tension resulting in differentiation and reduced immunosuppression^19^.

## Untargeted metabolomics points to respiration as a central mechanism

It has been previously reported that metabolite depletions unique to either the chondrogenic or osteogenic differentiation of MSCs can be identified by mass spectrometry^20^. Importantly, it has been shown that these metabolites could, by themselves, induce targeted differentiation^20^. We have developed this methodology to identify metabolites involved in MSC immunosuppression. However, a key challenge with studying basic MSC mechanisms and with elucidating MSC fate and function, is that MSC populations are highly heterogenous^21^. Thus, here, we use both an enriched skeletal (Stro-1^+^) MSC population^22^ and an unselected (total) commercial bone marrow-derived skeletal MSC population.

Total and Stro-1^+^ MSCs were cultured on SQ and on flat controls, and their metabolites were isolated at days 7 and 28 of culture, for mass spectrometry analysis. We annotated over 200 metabolites that changed in abundance between both MSC populations grown on SQ vs flat surfaces (Fig. 2A and Fig. S2). To provide focus, we selected only metabolites classified as true identifications (class I according to the Metabolite Standards Initiative guidelines^23^), providing 18 hits common to both time points (Fig. 2B). Of these, four metabolites were depleted at both days 7 and 28 (adenine, citrate, L-glutamic acid and niacinamide) (Fig. 2, C and D), which are all involved in cellular respiration (Fig. 2E).

**Fig. 2.**
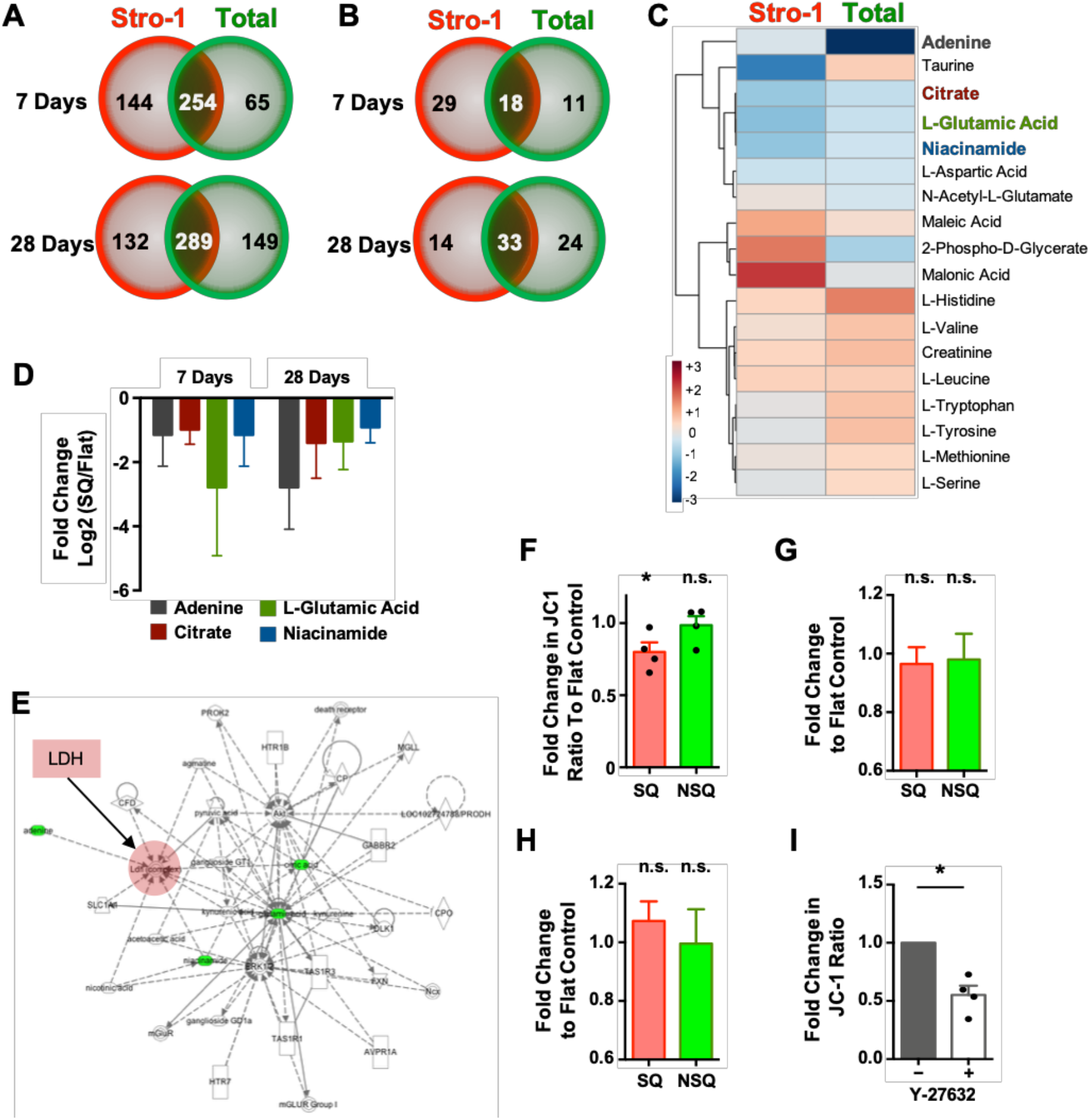
Metabolome analysis reveals nanotopography mediated changes in cellular respiration, independent of mitochondrial dynamics. (**A**) Stro-1^+^ (red) or total BM (green) MSCs were cultured on SQ or flat surfaces for 7 or 28 days, and the number of metabolites specific or common to both cell types were enumerated. Common metabolites with a confidence value of 10 were identified using IDEOM software in both MSC populations (**B**), and the heat map shows the distribution of these at day 7 of culture (**C**). (**D**) Fold change in selected metabolite concentrations. Data in (A-D) show mean of 6 nanotopographies per condition. (**E**) Biochemical network analysis of metabolite changes in MSCs cultured on flat versus SQ. (**F**) Changes in mitochondrial activity were measured using JC-1 staining, in MSCs cultured on SQ or NSQ nanotopographies relative to flat control. (**G** and **H**) Mitochondrial mass (Mitotracker Green) and superoxide generation (Mitosox Red) were also evaluated by flow cytometry. (**I**) Stro-1^+^ MSCs were cultured on flat and SQ nanotopographies for 14 days in the presence (hatched bars) or absence (open bars) of the ROCK inhibitor, Y27632, and changes in mitochondrial activity measured using JC-1 staining. Means ± SEM in F and G and number of donors (*N*) are shown for each condition. Means ± S.D. in D, G and H. \**p* < 0.05; n.s., non-significant.

There is a growing body of literature on pluripotent stem cells, which demonstrates that the Warburg effect^24^ supports the maintenance of their pluripotency^25,26^. The Warburg effect is a mechanism employed by cancer cells that involves a shift from oxidative phosphorylation in the mitochondria to oxidative glycolysis in the cytoplasm^24^. This metabolic shift is perhaps counterintuitive in fast-growing cancer and stem cells, as oxidative glycolysis is less efficient than oxidative phosphorylation is, in terms of adenosine triphosphate (ATP) production, which is the main source of cellular energy. However, this is mainly a problem for cells when glucose is deficient; in the body and in cell culture, glucose is in ready supply^27^.

Proliferating cells, in fact, might benefit from oxidative glycolysis by maintaining carbon rather than by releasing it as CO2, and from the increased metabolic supply of niacinamide adenine dinucleotide (NADH) and niacinamide adenine dinucleotide phosphate (NADPH). Both carbon supply and NADH/NADPH activity are important for amino acid, lipid and nucleotide biosynthesis, all of which are essential for generating new cells^27,28^. It is thus notable that the metabolites we have identified – adenine, citrate, L-glutamic acid and niacinamide – are all involved in cytoplasmic NADH/NADPH pathways^27,28^.

To determine if mitochondrial function is reduced in the immunosuppressive MSCs cultured on the SQ nanotopography, mitochondrial activity was assayed using JC1 staining and quantified by flow cytometry (Fig. S3). This showed that mitochondrial activity was reduced in MSCs cultured on SQ after 14 days of culture, as compared to MSCs cultured on the flat control (Fig. 2F). It is noteworthy that the measurement of mitochondrial mass, using mitotracker green, and of mitochondrial function, using superoxide generation with Mitosox red, demonstrated that MSCs cultured on the SQ or flat surfaces had comparable mitochondrial physiology (Fig. 2, G and H). These results suggest that in the presence of the SQ topography, mitochondrial activity is reduced but that function itself is not impaired.

We also inhibited ROCK signalling (via Y27632) in MSCs to determine if their mitochondrial activity, as measured by JC-1 expression, was affected by intracellular tension. Indeed, ROCK inhibition in MSCs cultured on flat control surfaces reduced JC-1 expression, indicating decreased mitochondrial activity (Fig. 2I) which pairs with the observation in Fig. 1, E-G that decreased intracellular tension increases immunosuppressive phenotype.

## Flux of heavy glucose shows increased oxidative glycolysis

To investigate changes in cellular respiration in MSCs grown on different nanotopographies, we used mass spectrometry to follow the breakdown and conversion of ^13^C-labelled glucose. Cells were cultured on SQ and flat control surfaces for 11 days in standard media followed by 3 days with ^13^C glucose-containing media. As illustrated in Fig. 3A, increased oxidative glycolysis results in increased lactate production. And indeed, MSCs cultured on SQ surfaces displayed increased glucose consumption and increased lactate production, relative to MSCs cultured on flat surfaces, while the mitochondrial tricarboxylic acid (TCA) cycle remained similar in MSCs cultured on either surface (Fig. 3B). Measurements of fluorescently labelled glucose uptake (Fig. 3C) and of extracellular lactate (Fig. 3D) confirmed the mass spectrometry data. Together, these results demonstrate that MSCs cultured on SQ, which display an enhanced immunosuppressive capability, subtly increase oxidative glycolysis, as is typically observed during the Warburg effect, compared to MSCs cultured on the flat control.

**Fig. 3.**
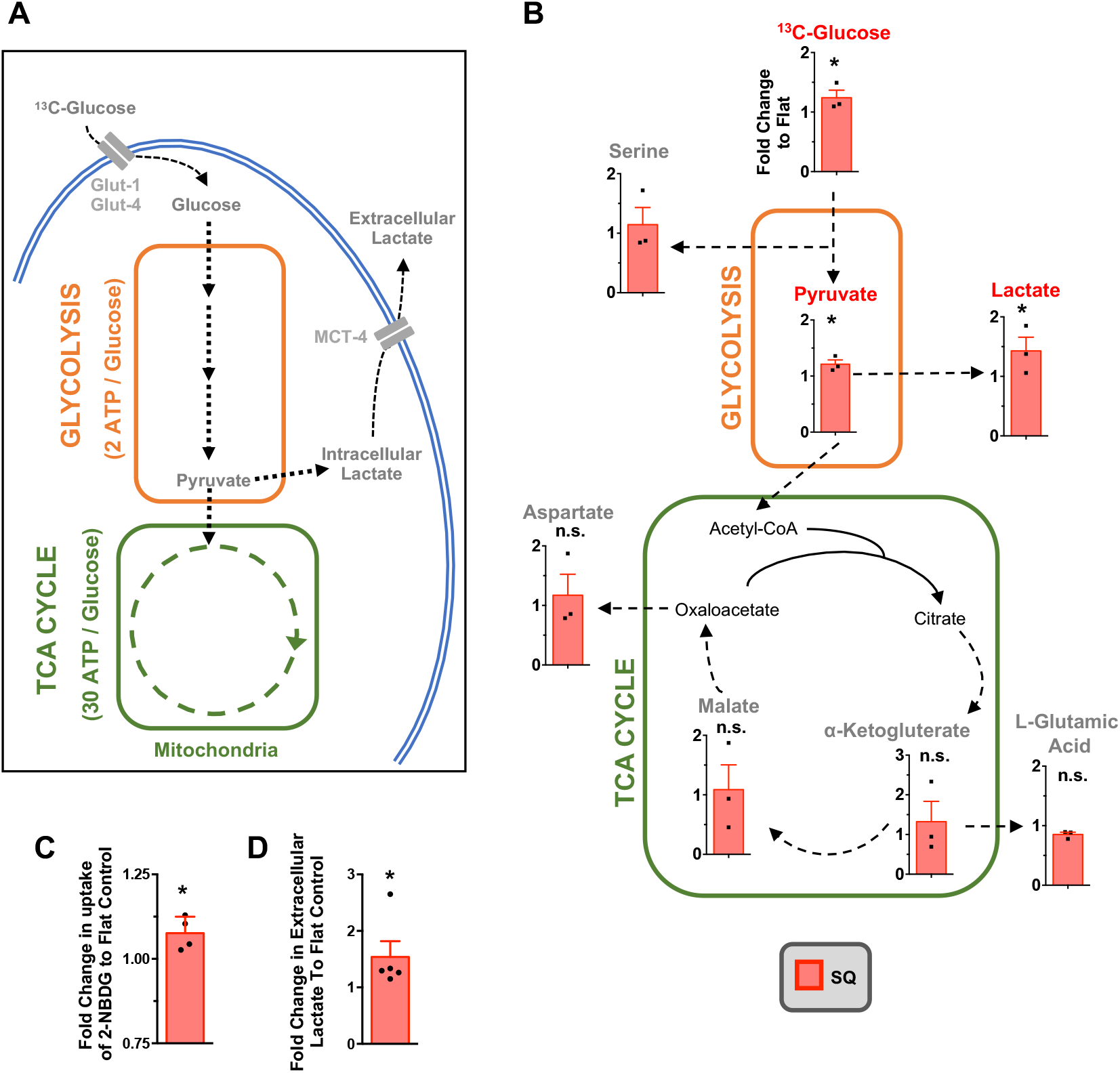
MSCs increase oxidative glycolysis on SQ nanotopographies, as revealed by [^13^C_6_]-glucose tracing. (**A**) Schematic of MSC respiration during culture. Changes to these pathways can be traced using heavy labelled [^13^C_6_]-glucose. (**B**) Stro-1^+^ MSCs were cultured for 14 days on nanotopographies in the presence of [^13^C_6_]-glucose for 72 hours. LC-MS was then used to measure the conversion and abundance of [^13^C_6_]-labelled metabolites in the glycolysis and TCA cycle pathways. Graphs show a fold change in [^13^C_6_]-labelled metabolites in MSCs cultured on SQ relative to flat nanotopographies. (**C**) MSCs were cultured for 14 days on SQ or flat surfaces, and glucose uptake was measured using 2-NBDG (a fluorescent glucose analogue) by flow cytometry. (**D**) Cell culture supernatants were collected from MSCs grown for 14 days on flat or SQ nanotopographies, and extracellular secreted lactate was quantified. Means ± SEM and number of donors (*N*) are shown for each condition. \**p* ≤ 0.05; n.s., nonsignificant.

No difference in glucose uptake, nor in oxidative glycolysis, was observed in MSCs cultured on the NSQ surface with ^13^C-labelled glucose, as compared to controls, but oxidative phosphorylation was increased, as indicated by enhanced L-glutamic acid production (Fig. S4). This finding supports the prevailing view that differentiation is energetically demanding for stem cells, and thus causes increased ATP production through increased oxidative phosphorylation^29–32^. It is further noteworthy that osteoblasts, as programmed from MSCs by the NSQ surface, are slow growing cells^19,33,34^. This highlights the need reduce oxidative phosphorylation for stem cell expansion.

## MSC immunosuppressive capacity enhanced by decoupling oxidative phosphorylation

The central aim of this study was to demonstrate that nanotopography can be used to identify pathways that can be exploited to maintain the immunomodulatory capabilities of MSCs in culture for a prolonged period of time. Our metabolomic analysis indicates that increased oxidative glycolysis could be key to achieving this aim. We hypothesized that by decoupling mitochondrial activity, to force cellular respiration to shift from oxidative phosphorylation to oxidative glycolysis, we could promote the MSC immunosuppressive phenotype. To test this, we used 2,4-dinitrophenol (DNP), an ionophore that dissipates proton gradients across mitochondria, preventing the proton motive force that produces ATP-related energy thus driving glycolysis^35^. MSCs were cultured on flat surfaces in the presence or absence of 0.5 mM DNP. We observed an increase in the immunosuppressive phenotype of DNP treated MSCs relative to untreated controls (Fig. 4A). One mechanism of MSC-mediated immunosuppression is the upregulation and release of indoleamine 2,3-dioxygenase (IDO), following MSC exposure to interferon-gamma (IFNγ), which is produced by activated T cells. IDO expression limits T cell proliferation through the degradation of extracellular tryptophan (a key amino acid required by T cells during proliferation), and is thus associated with immunomodulation^36^. After priming MSCs with 100ng/ml IFNγ overnight, we observed a similar upregulation of *IDO1* expression in both control and DNP-treated MSC populations (Fig. 4B). This demonstrates that DNP-treated MSCs which are utilising oxidative glycolysis for energy production can still respond to exogenous T cell stimuli in the form of IFNγ. The addition of DNP also promoted the retention of MSC markers, CD44, CD90 and CD166^37^ after 14 days in culture over that seen in untreated control cells (Fig. 4C). This shows that MSCs undergoing oxidative glycolysis retain markers of multipotency, as well as their immunosuppressive phenotype.

**Fig. 4.**
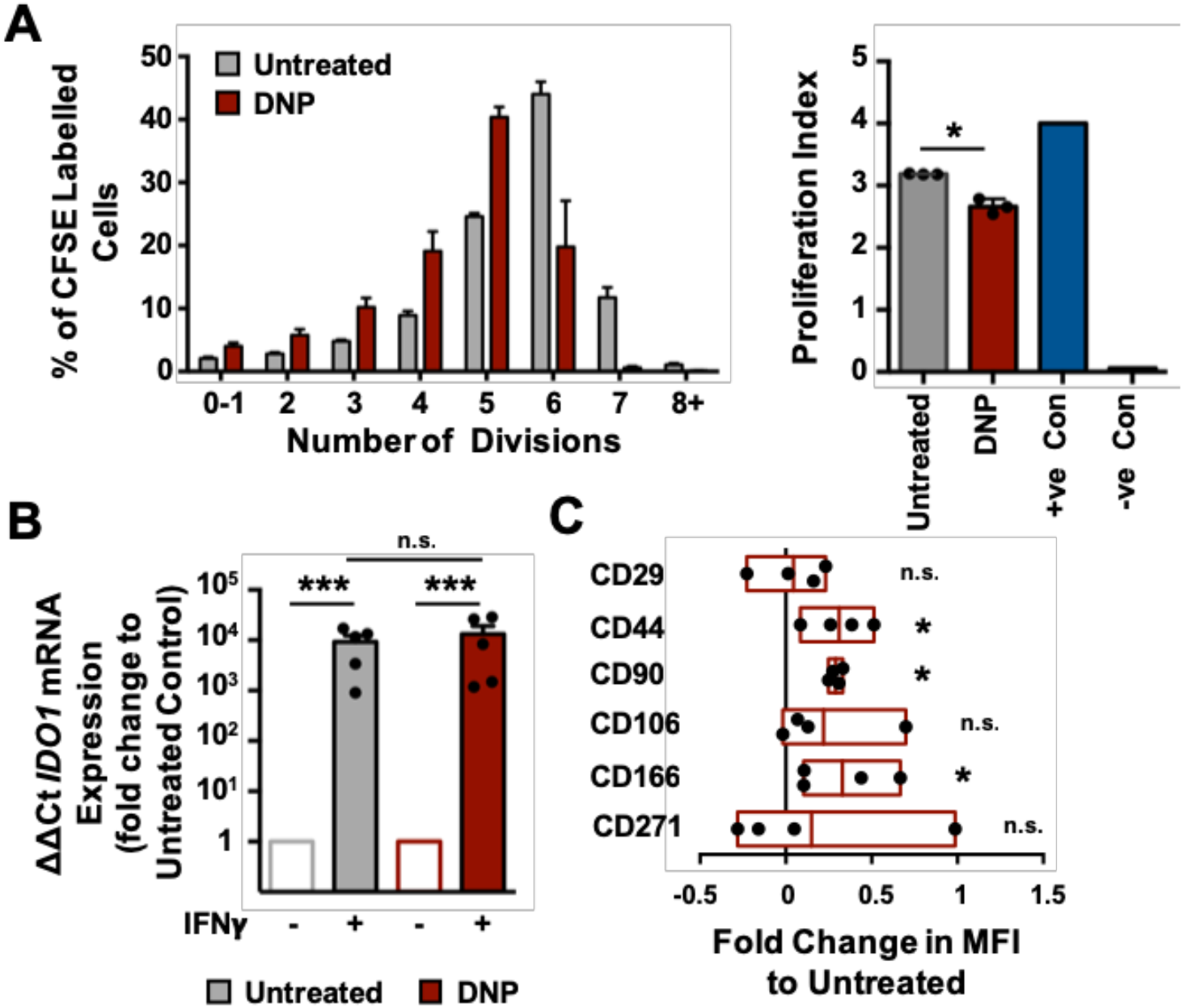
Uncoupling oxidative phosphorylation increases MSC immunosuppression. (**A**) Stro-1^+^ MSCs were cultured in the presence or absence of DNP for 14 days, then co-cultured with CFSE-labelled, IL-2 and PHA stimulated PBMCs for a further 5 days. Proliferation was assessed by flow cytometry. Data in left graph is a representative experiment (n=4 replicates per group; mean ± S.D.); data in right graph shows the proliferation index of 3 independent experiments. (**B**) Following culture with or without DNP, MSCs were challenged with IFN-γ for 24 hours and IDO1 expression was evaluated by qPCR. (**C**) Fold change in MSC surface marker expression following DNP treatment, relative to untreated controls, as assessed by flow cytometry. Graph shows mean with high and low values. Means ± SEM and number of donors (*N*) are shown for each condition. \*\*\**p* ≤ 0.0001; \**p* < 0.05; n.s., non-significant.

## Bioactive metabolites can enhance immunosuppressive capacity and growth

From our results, we hypothesised that the addition of individual metabolites identified in Fig. 2, A-D, namely adenine, citrate, niacinamide and L-glutamic acid, that were seen to link to respiration (Fig. 2E), might stimulate the MSC immunomodulatory phenotype. Indeed, when MSCs were treated with either adenine or L-glutamic acid, this significantly increased immunosuppression (adenine p = 0.0079; L-glutamic acid p = 0.0079) as revealed by coculture with CFSE labelled T cells (Fig. 5A). This agrees with our emerging hypothesis that MSC phenotype is related to the need to build new cells. Adenine is a purine nucleobase used for DNA synthesis in proliferating cells and is also a component of DNA and RNA and it also forms part of the NADH/NADPH dinucleotide (as well as ATP and flavin adenosine dinucleotide (FAD)). L-glutamic acid is an amino acid used in protein production. Thus, both directly fuel production of new cell components.

**Fig. 5.**
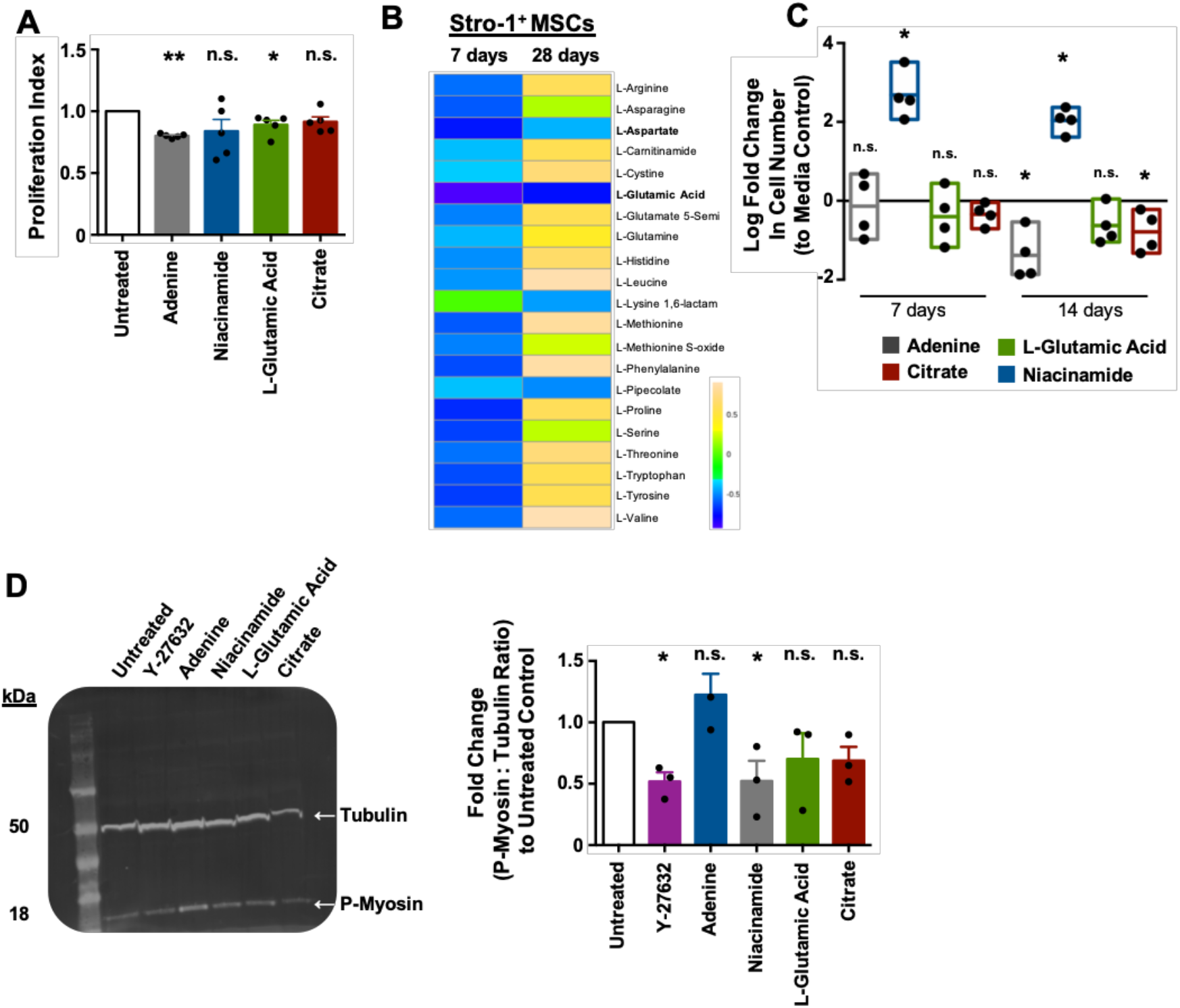
Addition of defined metabolite influences MSCs immunosuppressive capability. (**A**) Stro-1^+^ MSCs were cultured in the presence of metabolites for 14 days, followed by co-culture with CFSE labelled, IL-2 and PHA stimulated PBMCs for a further 5 days. Graph shows the proliferative index of T cells normalised to untreated controls and is representative of 2 independent experiments (n=4 topographies per group, mean ± S.D). (**B**) Changes in amino acid synthesis in Stro-1^+^ MSCs grown on SQ versus flat nanotopographies for 7 or 28 days. At both time points, L-glutamic acid and L-aspartate were depleted (n=6 topographies per group). (**C**) Stro-1^+^ MSCs were cultured with selected metabolites and fold change in total cell number was measured by flow cytometry relative to untreated controls. Graph shows mean with high and low values. (**D**) Stro-1^+^ MSCs were cultured with selected metabolites for 14 days or ROCK inhibitor Y-27632 for 7 days. Levels of phospho-myosin (18kDa) relative to β-tubulin (50kDa) was quantified by western blotting, Blot is representative of 3 independent donors. Graph on right shows quantitative changes in phospho-myosin expression normalised to β-tubulin. Means ± SEM and number of donors (*N*) are shown for each condition. \**p* < 0.05; n.s., non-significant.

Non-essential amino acid anabolism can be split into three categories^38,39^. In the first two categories, amino acid biosynthesis is TCA-cycle independent, with amino acids being derived from the pentose phosphate pathway and from glycolysis. Amino acids generated in this way include alanine, histidine, isoleucine, leucine, phenylalanine, serine, tryptophan, tyrosine and valine^38,39^. The third category consists of TCA cycle-derived amino acids, such as lysine, methionine and threonine, which are derived from canonical aspartate, and also arginine, glutamine and proline, which are derived from canonical glutamic acid (glutamate)^38,39^. Analysis of our untargeted metabolomic data from the Stro-1^+^ MSCs at days 7 and 28 showed that the canonical TCA cycle-derived amino acids, L-glutamic acid and L-aspartate, are depleted in the Stro-1^+^ MSCs, even at day 28 (Fig. 5B). We therefore propose that in the immunosuppressive MSCs, L-glutamic acid and L-aspartate become depleted because the TCA cycle does not increase in balance with oxidative glycolysis. This is interesting as many of these amino acids are considered to be conditionally essential and at times of high growth are required from diet^38,39^. Thus, we propose that as stem cells rapidly grow and employ glycolysis, they become depleted of these amino acids. It is also noteworthy that at days 7 and 14 of culture, the addition of niacinamide significantly increased Stro-1^+^ MSC growth (Fig. 5C). This again, illustrates that the identified metabolites are involved in new cell production.

This data allows us to postulate that cellular respiration links to cellular tension which influences MSCs immunosuppressive potential. In order to investigate this concept, Stro-1^+^ MSCs were cultured in the presence of metabolites identified in Fig. 2 for 14 days or Y27632 for 7 days. Western blot analysis of phospho-myosin revealed that addition of exogenous niacinamide acts to reduce intracellular tension in MSCs, comparable to addition of Y27632 (niacinamide p = 0.0277; Y27632 p = 0.0267) (Fig. 5D).

## Discussion

In this study, we set out to investigate whether nanotopography can enhance the immunomodulatory capacity of MSCs through increased oxidative glycolysis. A growing number of reports indicate that oxidative phosphorylation increases in MSCs, as well as in other stem cells, including pluripotent^29^ and hair follicle^30^ stem cells, as they undergo differentiation^32,40,41^, and that MSCs in standard culture are more glycolytic^32,40,41^. However, when MSCs are cultured on standard flat tissue culture plastics, they undergo phenotypical drift and lose certain characteristics, such as their immunosuppressive capacity, as shown in Fig. 1, C and D. In this study, we use the SQ nanotopography to regulate MSCs and to maintain their stem cell phenotype. This allows us to definitively show that shifting MSC respiration towards oxidative glycolysis is key to maintaining their immunosuppressive phenotype. Our data thus highlight the importance of properly controlling MSC experiments to correctly interpret data on cell function.

MSCs were first identified as precursors of mechanocytes^42^, now known as fibroblasts, and as fibroblast colony forming units^43^, due to the similarity in their appearance to fibroblasts and their ability to form stromal tissues. It is now emerging that subtle differences in intracellular tension exist between MSCs and fibroblasts, which provide a basis for understanding MSC behaviour in culture: maintained MSC phenotype (which has regenerative and therapeutic potential) versus phenotypic drift towards a fibroblast-like state^19^. MSCs have lower intracellular tension compared to fibroblasts^44^, and we demonstrate here that immunosuppression depends on this lowered cytoskeletal tension.

*In vivo,* in the bone marrow niche, hypoxia is likely to maintain enhanced glycolysis in MSCs via hypoxia inducible factor 1 (HIF-1α)^45,46^. HIF-1α promotes the expression of pyruvate dehydrogenase kinase, which prevents pyruvate from entering the TCA cycle, thereby inhibiting mitochondrial respiration^45,46^. Our analysis of RNAseq data, obtained from Stro-1^+^ MSCs cultured on SQ vs flat nanotopographies after 24 hours of culture, showed oxidative phosphorylation to be significantly repressed, but only a limited regulation of HIF1α signalling was observed (Fig. S5). This supports the hypothesis that in normoxia, on the SQ nanotopography, oxidative glycolysis is activated in MSCs through changes in cytoskeletal tension rather than via a hypoxic mechanism.

We propose that the tension-based mechanism that activates oxidative glycolysis can be exploited to support MSC growth in standard culture conditions, through the addition of specific metabolites that support cell growth, namely adenosine, L-glutamic acid, citrate and niacinamide. We propose that these metabolites are required for MSC growth, and for the maintenance of the MSC phenotype, by providing the precursors of anabolic co-factors and conditionally essential amino acids. Their provision could thus support enhanced MSC growth in the absence of oxidative phosphorylation, which is important because oxidative phosphorylation is associated with MSC differentiation^32,40,41^. The implications of these findings are important for the cell therapy field because they could be applied to support the large-scale expansion of MSCs for therapeutic application. Our results also demonstrate the value of material-based tools for understanding stem cell function and fate, and of their broader potential application across other biological systems and tissues.

## Acknowledgements

We thank Carol-Anne Smith for technical assistance. This work was supported by BBSRC funded grants BB/N018419/1, BB/K011235/1 and BB/L021072/1.

## Author Contributions

E.A.R., L-A.T., M.S-S. and M.J.D conceived and designed the analysis. E.A.R. performed the experimental work. L-A.T. performed the untargeted metabolomic experimental work. A.S., P.R. and N.G manufactured and provided polycarbonate nanotopographies. K.V.B and G.B performed metabolomic profiling and preliminary analysis of untargeted and ^13^C labelled cell extracts. J.A.W. and R.O.C.O provided Stro-1^+^ cells. E.A.R. and M.J.D. wrote the manuscript. L-A.T., G.B., M.S-S., J.M. and R.O.C.O. revised the manuscript and were involved in the discussion of the work.

## Competing Interests

Authors declare no competing interests.

## Notes

http://dx.doi.org/10.5525/gla.researchdata.778

