## Supplemental Figures, materials and methods for "Nanotopography reveals metabolites that maintain the immunosuppressive phenotype of mesenchymal stem cells"

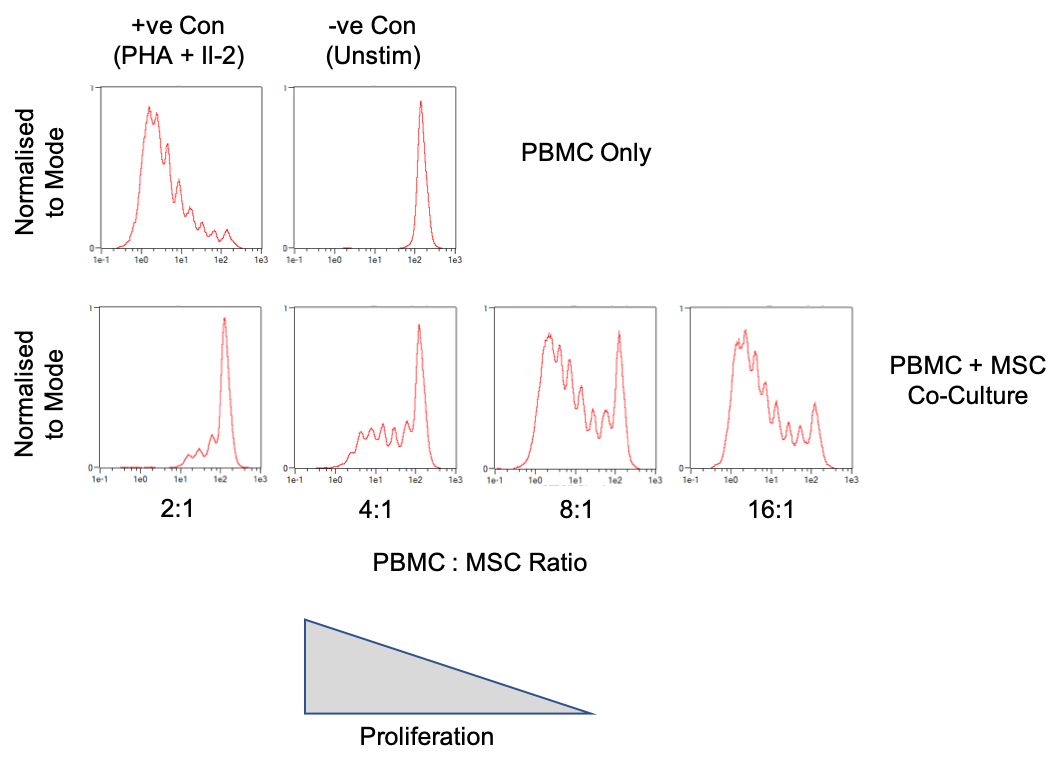

**Fig. S1. CFSE immunosuppression assay.** PBMCs were isolated from peripheral blood, labelled with CFSE, and stimulated to proliferate with 5μg/ml PHA-P and 100U/ml IL-2. PBMCs were then added to MSCs at defined ratios and co-cultured for 5 days. CFSE dilution was assessed by flow cytometry. Positive control was stimulated PBMCs on their own; negative control was CFSE-labelled PBMCs in the absence of PHA-P and IL-2.

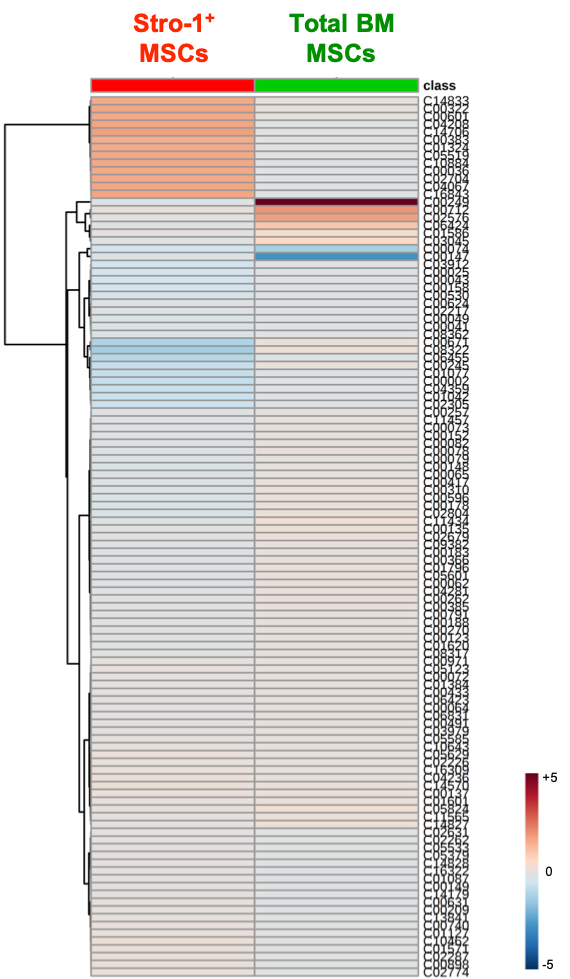

**Fig. S2. Changes in metabolic profiles of MSCs on topographies.** Stro-1^+^ (red) or total BM (green) MSCs were cultured on SQ or flat surfaces for 7 days, and the metabolic profile of MSCs was determined by analysis using IPA software. Heatmap is the mean of 6 individual topographies.

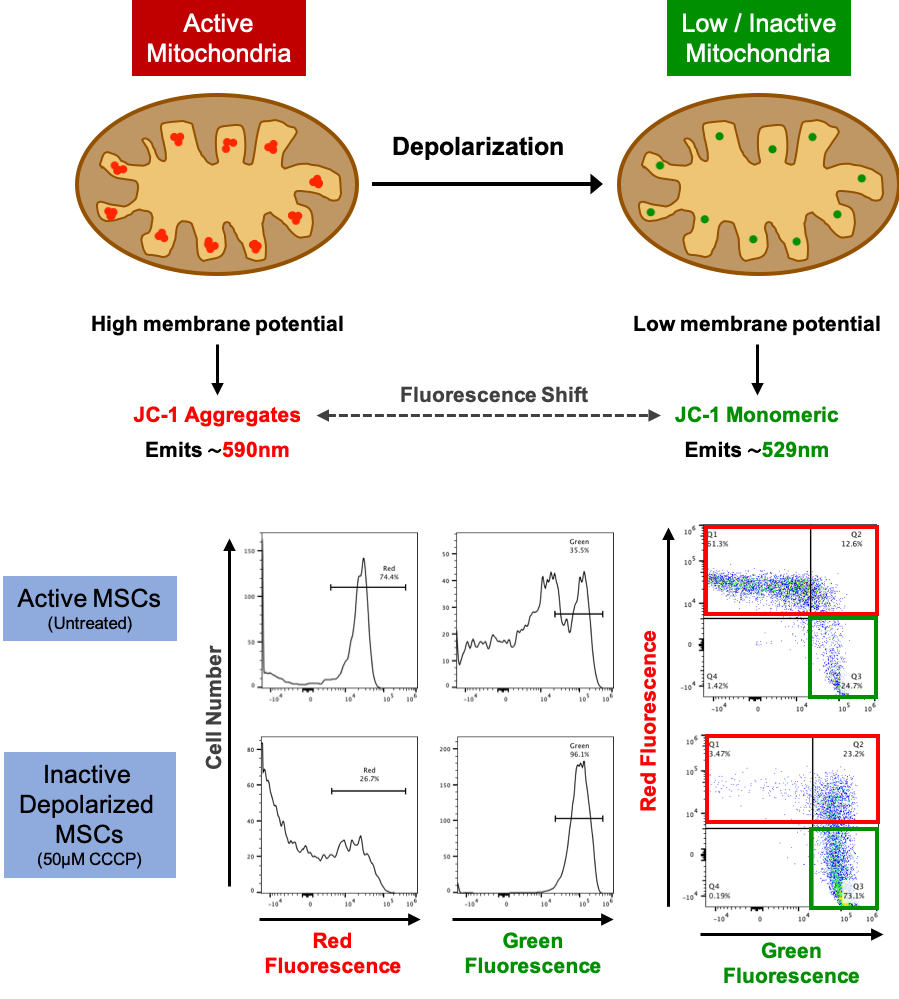
**Fig. S3. Mitochondrial activity measured using JC-1.** MSCs were labelled with 2 μM JC-1 for 30mins before being detached with trypsin and their fluorescence quantified by flow cytometry. Active mitochondria accumulate red fluorescent dye, whereas depolarised, inactive mitochondria remain green. Depolarisation can be induced by treating MSCs with 50 μM of the protonophore CCCP. Quantification of red and green fluorescence by flow cytometry provides a functional readout of mitochondrial activity. The JC-1 ratio allows mitochondrial function to be compared between individual experiments and donors.

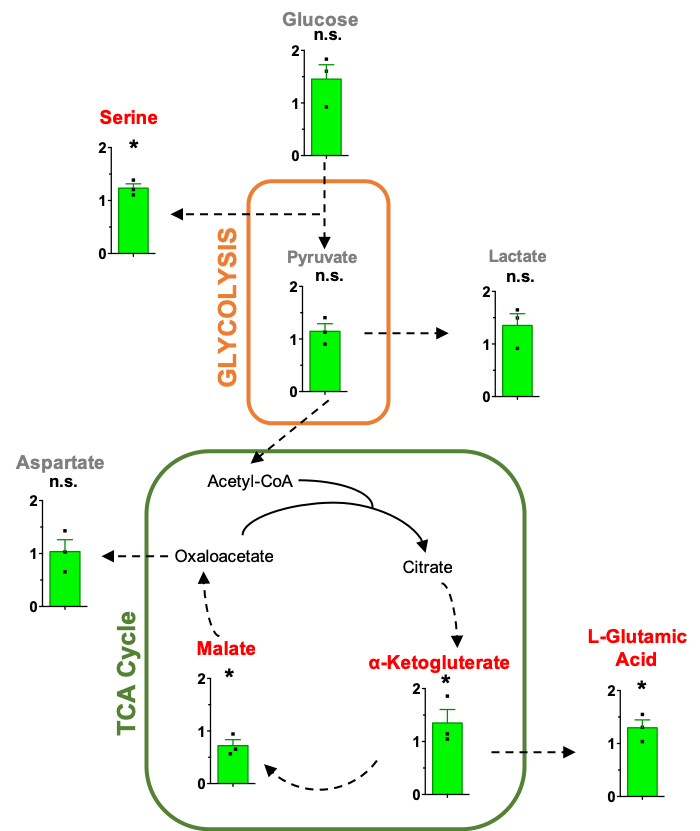

**Fig. S4. Metabolic tracing of [^13^C_6_]-glucose in MSCs cultured on NSQ nanotopography.** Stro-1^+^ MSCs were cultured on NSQ or flat nanotopographies for 11 days, followed by a further 3 days in the presence of [^13^C_6_]-glucose. LC-MS was used to measure the conversion and abundance of [^13^C_6_]-labelled metabolites. Graphs show the fold change in [^13^C_6_]-labelled metabolites in MSCs cultured on NSQ relative to flat nanotopographies. The results show increase in mitochondrial respiration as indicated by increased ^13^C incorporation in ketoglutarate and malate. (n=3 independent experiments; each point is the mean of 4 topographies per group; mean ± SEM, ^*^P<0.05).

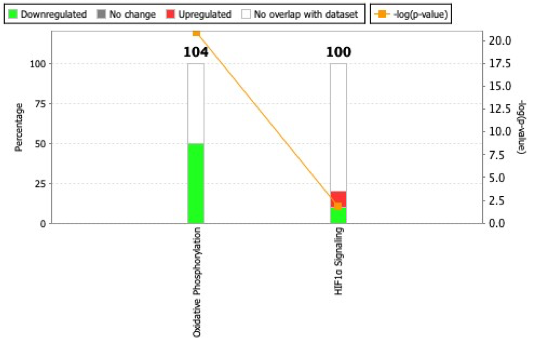

**Fig. S5. RNAseq reveals changes to oxidative phosphorylation.** Stro-1^+^ MSCs were cultured on SQ or flat nanotopographies for 24 hours before RNA was harvested and analysed by next generation sequencing. Changes to RNA species involved in oxidative phosphorylation and HIF-1α pathways were evaluated using Ingenuity Pathway Analysis software. N=3 material replicates per group.

**Materials and Methods**

Materials fabrication. Two master substrates were fabricated on silicon coated with 100 nm PMMA (Elvacite 2041, Lucite International) by electron beam lithography to generate arrays of 120 nm diameter pits with 300 nm centre-centre spacing and 100 nm depth, corresponding to the SQ nanotopography, or arrays with an additional ± 50 nm offsetting of pits for NSQ. The silicon substrates were exposed to a 50 kV electron beam, developed in 1:3 MIBK:IPA for 30 s, rinsed in IPA and dried in a nitrogen stream. Nickel dies were made from the patterned resists and 50 nm of Ni-V was sputtered coated. Electroplating was carried out to a thickness of approx. 300 μm (outsourced to DVDNorden, Denmark). These nickel shims were cleaned with chloroform for 10-15 min in an ultrasonic bath, subjected to further rinses in acetone and IPA, and dried once more in gaseous nitrogen. Polycarbonate (Makrolon^®^ OD2015) substrates were generated by injection moulding using an Engle Victory 28 hydraulic injection moulder. The required nickel inlay corresponding to a SQ arrangement or NSQ arrangement was inserted prior to production. Heating to 180°C melted the polycarbonate. A clamping force of 250 kN was used to imprint onto the surface of the polycarbonate, with the final dimensions of each substrate being 24 mm x 24 mm. The temperature was allowed to drop to 70°C before separation of the press and polymer. Unpatterned (flat) polycarbonate substrates were injection moulded against planar shims and used as controls.

MSC isolation and culture. Stro-1^+^ MSCs were isolated from bone marrow obtained from hematologically normal patients undergoing routine knee/hip replacement, as previously described^1^. Commercial BM MSC were purchased from PromoCell (PromoCell GMBH, Germany). In all experiments, cells from multiple donors were used, as outlined in the Fig. legends and summarised in Supplementary Table 2. MSCs were routinely cultured in basal media (comprising DMEM supplemented with 10% (v/v) Foetal Calf Serum, 2.5 mM L-glutamine, 80 U/ml penicillin, 0.11 mM streptomycin, 1x non-essential amino acids, 1 mM sodium pyruvate (all Sigma-Aldrich) and 0.2 µg/ml Fungizone amphotericin B (Gibco)) and used for experiments at passages 2-4. Cells were seeded on nanotopographies at 1000 cells/cm^2^ using seeding devices as previously described^1^ and cultured for 14 days. Basal culture media was replaced every 2-3 days.

Influencing MSC Respiration in vitro. MSCs were forced to into a glycolytic state by culturing them for 14 days in the presence of 0.5 mM 2,4-Dinitrophenol (DNP), a mitochondrial decoupling protonophore. Media was replaced every 2-3 days. To assess the impact of extracellular metabolites on cell function, MSCs (1000 cells/cm^2^) were cultured in basal media supplemented with either 5 mM adenine, 5 mM Citrate, 10 mM Niacinamide or 25 mM L-Glutamic Acid. Media was replaced every 2-3 days. To assess the effect of cytoskeletal tension on respiration, MSCs were cultured for 7 or 14 days in basal media containing 10 µM of the ROCK inhibitor Y27632 (Abcam). Media was changed every two days.

Flow Cytometry. To phenotype MSCs after culture, cells were washed once with PBS and detached using Accutase (ThermoFisher). Cells were stained on ice for 30-45 mins using antibodies outlined in Supplementary Table 1 in flow cytometry buffer (PBS supplemented with 0.5% BSA and 0.5 mM EDTA). Cells were then washed twice with PBS and analysed using an Attune NXT flow cytometer (ThermoFisher).

**Table S1.** Reagents used for flow cytometry.

| Antigen | Clone | Isotype | Fluorochrome | Source |
| --- | --- | --- | --- | --- |
| CD29 | TS2/16 | mIgG1 | FITC | eBioscience |
| CD44 | IM7 | Rat IgG2b | PE-Cy7 | eBioscience |
| CD90 | eBio5E10 | mIgG1 | PerCP-eFluor710 | eBioscience |
| CD106 | STA | mIgG1 | PE | eBioscience |
| CD166 | 3A6 | mIgG1 | PerCP-eFluor710 | eBioscience |
| CD271 | REA844 | REA | PE-Vio770 | Miltenyi Biotech |

| Isotype Control | Clone | Fluorochrome | Source |
| --- | --- | --- | --- |
| Mouse IgG1k | P3.6.2.8.1 | FITC | eBioscience |
| Rat IgG2b | eB149/10H5 | PE-Cy7 | eBioscience |
| Mouse IgG1k | P3.6.2.8.1 | PerCP-eFluor710 | eBioscience |
| Mouse IgG1k | P3.6.2.8.1 | PE | eBioscience |
| Recomb hum IgG1 | REA293 | PE-Vio770 | Miltenyi Biotech |

Mitochondrial function was quantified using the membrane potential dye JC-1 (ThermoFisher). Cells were cultured in basal media containing 2 μM JC-1 for 30 mins at 37°C, then detached and analysed by flow cytometry. Cells treated with JC-1 and 50 μM carbonyl cyanide 3-chlorophenylhydrazone (CCCP, SigmaAldrich) were used as a positive control for mitochondrial depolarisation. Mitochondrial mass was measured by incubating cells with 100 nM MitoTracker Green (ThermoFisher) in serum free media for 30 mins at 37°C. MSCs were then washed, detached and mass analysed. Mitochondrial superoxide generation was quantified by incubating MSCs with 5 μM MitoSOX Red in HBSS for 10 min at 37°C, washing, detaching and measuring intracellular fluorescence.

Glucose uptake was measured by culturing cells in glucose-free basal media for 2 hours before adding the fluorescently tagged glucose analogue 2-NBDG (2-(N-(7-Nitrobenz-2-oxa-1,3-diazol-4-yl) Amino-2-Deoxyglucose) for 60 mins at 37°C (ThermoFisher). MSCs were washed, detached and uptake of glucose measured by fluorescence incorporation.

Flow cytometry files were analysed using FlowJo software (version 10.5.3, FlowJo LLC, USA). In all experiments, a minimum of 5000 cells per sample were analysed to ensure statistical significance.

Immunostaining. Topographies were washed twice with PBS, and cells fixed in a solution of 4% paraformaldehyde for 15 mins at 37°C. Fixed cells were permeabilised for 5 mins at 4°C in perm buffer (50 mM NaCl, 30 mM sucrose, 3 mM MgCl_2_·6H_2_O, 20 mM HEPES, 0.5% v/v Triton X-100 in PBS, pH 7.2) followed by blocking for 60 mins at RT in blocking buffer (1% BSA (v/v) in PBS). Cells were stained with rhodamine tagged phalloidin (ThermoFisher) for 1 hour in blocking buffer at RT. Nanotopographies were washed for 6x 5 mins with wash buffer (PBS + 0.5% (v/v) Tween^©^ 20 (SigmaAldrich)) and mounted with immunomount solution containing DAPI to stain nuclei (Vectorshield, Vector Laboratories). Cells were imaged using an Axiophot fluorescence microscope using x10 and x40 Neofluor objectives (all Zeiss), and captured with an Evolution QEi digital camera (Media Cybernetics, Rockville, USA) using QCapture software (QCapture Suite Plus, version 3.1.3.10, Teledyne QImaging, Surrey, BC, Canada). Actin expression was quantified using ImageJ software (version 1.52a) and normalised to nuclei count per field. 15 fields per topography were analysed.

Cell Culture Lactate Measurements. MSCs were cultured on nanotopographies for 11 days in normal basal media. Cells were then washed twice with serum free DMEM and cultured for the final 72 hours in basal media where normal FCS was substituted with 10% (v/v) dialysed FCS (ThermoFisher). Cell culture supernatants were collected, aliquoted and frozen at -80°C. Lactate levels in supernatants were quantified using the Lactate-Glo chemiluminescence assay (Promega), and luminescence measured using a Pherastar FS plate reader (BMG Labtech).

Gene Expression. After culture, RNA was isolated from MSCs by lysing cells in 350 ul Trizol reagent (Life Technologies). 200 ul/ml of chloroform (Sigma) was added per 1 ml of Trizol, mixed, centrifuged and the aqueous phase removed. Total RNA was extracted using the RNeasy extraction kit (Qiagen) according to the manufacturer’s protocol. Purified RNAs were quantified using a Nanodrop ND1000 spectrophotometer (Thermo Scientific) and cDNA synthesised with the Qiagen Quantitect reverse transcription kit, according to the manufacturer’s protocol. PCR amplification of target genes was performed using Quantifast SYBR green qPCR kit (Qiagen) with specific primers (Eurofins). PCR was quantified using the 2^-ΔΔCt^ method and amplification performed using an Applied Biosystems 7500 Real Time PCR System and the fold upregulation of genes was compared to untreated controls.

Immunosuppression Assay. Peripheral blood mononuclear cells (PBMC)s were purified from buffy coats by layering onto Ficoll Plus (GE Healthcare) density gradients as per the manufacturer’s instructions. PBMCs were labelled with 5 μM CellTrace CFSE (ThermoFisher) in PBS for 20mins at 37°C. Labelled PBMCs were counted and resuspended in stimulation media (basal media containing 5 μg/ml Phytohemagglutinin (PHA)-P (Sigma Aldrich) and 100 U/ml IL-2 (PeprotechEC)) to induce T lymphocyte proliferation. Stimulated CFSE labelled cells were added to MSCs at a 4:1 ratio (4 PBMC to 1 MSC) unless otherwise stated. Co-cultures were incubated for five days, before passing suspended cells through a 100 μm cell filter for analysis proliferation by CFSE dilution. Proliferation controls included in each assay were CFSE labelled PBMCs in stimulation media alone (positive control), and CFSE labelled PBMCs in basal media only (negative control). In order to compare independent assays, the proliferation index was calculated using the proliferation analysis plugin in FlowJo software (version 10.5.3, FlowJo LLC, USA).

Metabolomic Analysis. Whole cell metabolomic analysis was performed on cell lysates isolated from MSC cultured on nanotopographies for 7 or 28 days. Nanotopographies were washed with ice-cold PBS, and cells lysed in extraction buffer (PBS/methanol/chloroform at 1:3:1 Ratio) for 60 mins at 4°C with constant agitation. Lysates from duplicate nanotopographies were pooled as one sample, and extracts were transferred to cold eppendorfs and spun at 13,000 g at 4°C for 5 mins to remove debris. Cleared extracts were used for hydrophilic interaction liquid chromatography-mass spectrometry analysis (UltiMate 3000 RSLC, (ThermoFisher), with a 150 x 4.6 mm ZIC-pHILIC column running at 300 µl/min^-1^ and Orbitrap Exactive). Sample protein concentrations were measured by Nanodrop and used to standardise samples where required. A standard pipeline, consisting of XCMS (peak picking), MzMatch (filtering and grouping) and IDEOM (further filtering, post-processing and identification) was used to process the raw mass spectrometry data. Identified core metabolites were validated against a panel of unambiguous standards by mass and predicted retention time. Further putative identifications were generated by mass and predicted retention times. Means and standard errors of the mean were generated for every group of picked peaks and the resulting metabolomics data were uploaded to Ingenuity Pathway Analysis (IPA, Qiagen) software for metabolite pathway analysis. Heat maps of selected metabolites were generated using MetaboAnalyst software (version 4.0^2^).

^13^C_6_-Glucose Metabolomic Tracing. MSC were seeded on nanotopographies at 1000 cells/cm^2^ and allowed to grow for 11 days. Cells were washed and incubated in basal media comprising 50% normal glucose and 50% ^13^C_6_-Glucose (Cambridge Isotopes Ltd) for a further 3 days. Extractions were performed as above. LC-MS was performed as previously described^3^. Briefly, The LC-MS platform consisted of an Accela 600 HPLC system combined with an Exactive (Orbitrap) mass spectrometer (ThermoFisher). Two complementary columns were used; the zwitterionic ZIC-pHILLIC column (150 mm x 4.6 mm; 3.5 μm, Merck) and the reversed phase ACE C18-AR column (150 mm x 4.6 mm; 3.5 μm Hichrom) and in both cases sample volume was 10 μl at a flow rate of 0.3 ml/min. Eluted samples were then analysed by mass spectrometry.

Raw data from LCMS of ^13^C-labelled extracts was processed to generate a combined PeakML file as described previously^3^. Further analysis using mzMatch-ISO in R^4^ generated a PDF file containing chromatograms used to check peak-shape and retention time, and a tab-delineated file detailing peak height for each isopotologue, which was used to calculate percentage labelling.

Next-generation Sequencing (RNAseq). After culture, RNA was isolated from MSCs using the RNeasy extraction kit (Qiagen) following the manufacturer’s instructions. RNA concentrations were quantified using a Nanodrop ND1000 spectrophotometer (ThermoFisher) and samples were submitted to the Glasgow Polyomics facility (University of Glasgow). A Truseq^©^ stranded mRNA low throughput kit (Illumina) was used to process 136 ng of RNA per sample. A library of template molecules for sequencing was generated by conversion of mRNA (using poly-A selection, first and second strand cDNA synthesis, cleanup, adenylation of 3′ ends, ligation and cleaning of adaptors). PCR amplification was then performed (98°C for 30s, then 15 cycles of; 98°C for 10s, 60°C for 30s and 72°C for 30s followed by 72°C for 5 min). PCR products were cleaned using AMPure XP beads (Beckman Coulter). Sequencing of 3.2 pM libraries using a Nextseq 500 analyser (Illumina). Initial bioinformatics was performed using a BaseSpace^©^ next-generation sequencing platform (Illumina), with differential expression data being analysed with IPA software (Qiagen).

Western Blotting. MSCs were cultured in 25cm^2^ flasks in the presence or absence of metabolites or Y-27632 inhibitor. Cell were washed once with cold PBS, then lysed in 200ul RIPA buffer (150mM NaCl, 1% (v/v) NP-40, 0.1% (v/v) Sodium dodecyl sulfate in 50mM Tris HCl pH8) supplemented with protease and phosphatase inhibitors (Pierce Inhibitor Mini Tablets, ThermoFisher). Lysate protein concentration was measured (Pierce BCA Protein Assay Kit, ThermoFisher) and 20μg loaded per lane of a Bolt 4-12% Bis-Tris Gel using a XCell SureLock gel tank (both ThermoFisher). Protein was transferred onto Immobilon-FL PVDF membrane (Merck) and blocked for 1 h at RT with 5% (v/v) nonfat dry milk powder in Tris Buffered Saline (TBS). Membranes were probed overnight at 4°C with mouse anti-human phospho-myosin light chain 2 (Ser19, clone #3675, Cell Signalling Technology) and mouse anti-human β-tubulin (clone TUB 2.1, Sigma Aldrich) in TBS with 5% (v/v) BSA. Membranes were washed with TBST (TBS + 0.5% (v/v) Tween-20) for 6 x 5 min, and probed with IRDye 800CW goat anti-mouse IgG secondary antibody (Li-Cor) in TBS containing 5% (v/v) BSA for 1 h at RT. Membranes were washed for 6 x 5 min and imaged using an Odyssey Sa infrared imaging system with Image Studio 4.0 software (Li-Cor). Bands were quantified by densitometry using ImageJ software^5^.

Statistics. Unless stated, two-tailed unpaired T tests (Mann-Whitney) were performed where a direct comparison was assessed between two population groups using Prism software (GraphPad). A sample population of between 3-7 replicates was always used. Results are quoted as mean ± standard error of the mean or standard deviation, where appropriate. The probability values are quoted to an accuracy of 95%, 99% and 99.9% (**P* < 0.05, ***P < 0.01 and *P* ≤ 0.001 respectively). Supplementary Table 2 indicates sample numbers and replicates in each figure presented.

**Table S2.** Summary of cell donors used and experimental replicates.

| **Figure** | **Experiment** | **Number of Donors** | **Biological Replicates** | **Technical Replicates** |
| --- | --- | --- | --- | --- |
| 1c | Immunosuppression | 1 | 1 | 4 |
| 1d | Immunosuppression | 7 | 7 | 3-4 |
| 1e | Y27632 + Actin | 1 | 1 | 15 |
| If | Y27632 + Immunosuppression | 1 | 1 | 3 |

| 1g | Y27632 + Immunosuppression | 3 | 3 | 4 |
| --- | --- | --- | --- | --- |

| 2a, b, c | Metabolomics | 3 | 3 | 4 |
| --- | --- | --- | --- | --- |
| 2d | Metabolomics | 3 | 3 | 4 |
| 2f | JC1 | 4 | 4 | 3 |
| 2g | MitoTracker | 1 | 1 | 4 |
| 2h | MitoSOX | 1 | 1 | 4 |
| 2i | Y27632 + JC1 | 4 | 4 | 3-4 |
| 3b | 13C-Glucose | 3 | 3 | 3-4 |
| 3c | 2-NBDG | 4 | 4 | 3-4 |
| 3d | Lactate | 3 | 3 | 1-2 |

| 4a | DNP + Immunosuppression | 3 | 3 | 4 |
| --- | --- | --- | --- | --- |

| 4b | qPCR | 5 | 5 | 1 |
| --- | --- | --- | --- | --- |
| 4c | Phenotyping | 4 | 4 | 1 |
| 5a | Immunosuppression | 2 | 2 | 4 |
| 5b | Metabolomics | 3 | 3 | 4 |
| 5c | Metabolites and cell number | 4 | 4 | 1 |
| 5d | P-Myosin Western Blot | 3 | 3 | 1 |
